# Generation and characterization of a multi-functional panel of monoclonal antibodies for SARS-CoV-2 research and treatment

**DOI:** 10.1101/2023.11.08.566276

**Authors:** Lila D. Patterson, Benjamin D. Dubansky, Brooke H. Dubansky, Shannon Stone, Mukesh Kumar, Charles D. Rice

## Abstract

The Coronavirus disease 2019 (COVID19) pandemic caused by Severe Acute Respiratory Syndrome-Coronavirus-2 (SARS-CoV-2) is an ongoing threat to global public health. To this end, intense efforts are underway to develop reagents to aid in diagnostics, enhance preventative measures, and provide therapeutics for managing COVID-19. The recent emergence of SARS-CoV-2 Omicron variants with enhanced transmissibility, altered antigenicity, and significant escape of existing monoclonal antibodies and vaccines underlines the importance of the continued development of such agents. The SARS-CoV-2 spike protein and its receptor binding domain (RBD) are critical to viral attachment and host cell entry and are primary targets for antibodies elicited from both vaccination and natural infection. In this study, mice were immunized with two synthetic peptides (Pep 1 and Pep 2) within the RBD of the original Wuhan SARS-CoV-2, as well as the whole RBD as a recombinant protein (rRBD). Hybridomas were generated and a panel of three monoclonal antibodies, mAb CU-P1-1 against Pep 1, mAb CU-P2-20 against Pep 2, and mAb CU-28-24 against rRBD, were generated and further characterized. These mAbs were shown by ELISA to be specific for each immunogen/antigen. Monoclonal antibody CU-P1-1 has limited applicability other than in ELISA approaches and basic immunoblotting. Monoclonal antibody CU-P2-20 is shown to be favorable for ELISA, immunoblotting, and immunohistochemistry (IHC), however, not live virus neutralization. In contrast, mAb CU-28-24 is most effective at live virus neutralization as well as ELISA and IHC. Moreover, mAb CU-28-24 was active against rRBD proteins from Omicron variants B.2 and B.4/B5 as determined by ELISA, suggesting this mAb may neutralize live virus of these variants. Each of the immunoglobulin genes has been sequenced using Next Generation Sequencing, which allows the expression of respective recombinant proteins, thereby eliminating the need for long-term hybridoma maintenance. The synthetic peptides and hybridomas/mAbs are under the intellectual property management of the Clemson University Research Foundation, and the three CDRs have been submitted as an invention disclosure for further patenting and commercialization.

## INTRODUCTION

The Severe Acute Respiratory Syndrome-Coronavirus-2 (SARS-CoV-2) and associated severe pneumonia first appeared in the Wuhan Province of China in late 2019, and within weeks had spread globally, resulting in the COVID-19 pandemic [1]. Early epidemiological monitoring estimated the SARS-CoV-2 reproduction number (*R*_0_) of the parental (Wuhan) strain of SARS-CoV-2 between 2.2-2.9, with a doubling time of 5 days [2]. The *R*_0_value of SARS-CoV-2 indicates the virus is more transmissible than SARS-CoV, likely due to differences in the SARS-CoV-2 receptor binding domain that enhance binding affinity to the host receptor ACE2 [3]. The primary route of SARS-CoV-2 transmission is via respiratory droplets during unprotected close contact with an infected individual. Additionally, the occurrence of aerosol transmission is well-reported in crowded, poorly ventilated indoor settings and within hospitals, where aerosol-generating procedures such as respiratory therapies are common [3]. In contrast to severe SARS-CoV-2, asymptomatic and pre-symptomatic transmission is common and is thought to contribute to a significant number of infections [4].

Most COVID-19 infections do not require hospitalization, typically resolving within two weeks. Non-hospitalized patients with mild to moderate disease who are at risk of progression to severe disease may be prescribed one of the three antiviral medications: oral nirmatrelvir *(Paxlovid),* intravenous remdesivir *(Veklury),* and oral molnupiravir *(Lagevrio)*. Paxlovid is FDA-approved for such use, however, Veklurv and Lagevrio are only available under the Emergency Use Authorization (EUA) [5-9]. Antiviral medications are typically combined with the corticosteroid dexamethasone and immunomodulators (IL-6 inhibitors, JAK inhibitors) to reduce the inflammatory response.

Monoclonal antibodies (mAbs) targeting antigenic regions of viral surface proteins are highly valuable agents for the detection, treatment, and prevention of disease progression. The urgent need for such tools has prompted the rapid development of a vast number of monoclonal antibodies targeting SARS-CoV-2. In convalescent patients and in individuals post vaccination, most of the circulating IgG antibodies are against the Spike 1 (S1) epitopes outside of the receptor binding domain. While these antibodies predominate, they are typically not neutralizing antibodies [10]. In contrast, most SARS-CoV-2 neutralizing antibodies are generated against the RBD [11,12]. As of June 2022, five anti-SARS-CoV-2 monoclonal antibodies received EUA from the FDA for the treatment and prevention of COVID-19 in non-hospitalized patients at high risk for severe disease [13]. All monoclonal antibodies previously granted EUA are directed against the SARS-CoV-2 S1, including the RBD protein, inhibiting the interaction between the RBD and ACE2 and blocking viral entry and infection. Clinical trials showed SARS-CoV-2 neutralizing monoclonal antibodies to be safe and well tolerated, and early administration of was found to reduce the risk of disease progression by up to 80% [14]. When given to asymptomatic individuals, monoclonal antibody treatment early in the pandemic was found to reduce the incidence of symptomatic infection and was also shown to be effective in the prevention of disease or infection in those exposed to SARS-CoV-2 [14].

The continued development of monoclonal antibodies against SARS-CoV-2 RBD is essential for additional efforts to control the spread of COVID-19 and combat current and future variants. However, since the evolution of the Omicron variant, approved mAbs for COVID-19 therapy no longer demonstrate efficacy in mitigating the clinical progression of disease [15]. One mAb, Bebtelovimab, originally held promise as it neutralized the early Omicron variants. However, Bebtelovimab has become less effective due to the evolution of newer, more resistant variants [16]. Despite rapid advances in the development of therapeutics for SARS-CoV-2-related pathologies, there remains the need for monoclonal therapeutics targeting recent (and future) variants of concern.

To date, most efforts to develop mAb therapies for COVID-19 are focused on neutralizing antibodies [17], though mAbs have many other properties and applications. For example, an antibody may lack the ability to neutralize the SARS-CoV-2, but may well recognize epitopes revealed during ELISA applications, immunoblotting, and immunohistochemistry. Likewise, mAbs that have limited use in these areas may be strong neutralizing antibodies. In the study described herein, a simplified adaptation of current COVID-19 vaccines based on spike mRNA or expression of recombinant spike proteins were used to generate a panel of mAbs. These mAbs were used to test the hypothesis that immunogenic synthetic peptides (15 – 20 amino acids) can induce the production of high quality mAbs for multiple applications in SARS-CoV-2 research and treatments. Specific immunogenic synthetic peptides derived from the RBD region of the S proteins were conjugated to a large carrier protein (keyhole limpet hemocyanin; KLH). Mice were then immunized for the generation of polyclonal anti-sera and mAbs, followed by the comparison to antibodies generated. These were then tested for their ability to neutralize the original Wuhan strain of SARS-CoV-2 using plaque reduction neutralization test (PRNT) and recognize the RBD by ELISA, immunoblotting, and immunohistochemistry. One mAb, mAb CU-28-24, not only neutralizes the original Wuhan strain, but is also highly cross reactive against recent Omicron variants BA.2 and BA.4.5.

## MATERIALS AND METHODS

### Immunogen design

Synthetic peptides within two internal amino acid sequences (between 400 – 500) of the receptor binding domain (Arg319 – Phe541) of the original Wuhan strain of SARS-CoV-2 spike protein were designed from a full-length RBD cDNA sequence (accession # NC_045512.2) and conjugated with keyhole limpet hemocyanin (KLH) (Anaspec, Richmond CA) for immunizations and an unconjugated peptide for screening assays. Recombinant RBD (Arg319 – Phe541) of the same strain was obtained from Invitrogen (# RP 87678) and used as an immunogen without conjugation with KLH and used for screening assays. Sequences of the two synthetic peptides were chosen based on Hopp-Woods hydrophilicity profiles (https://web.expasy.org/protscale/), NIH-Ab-designer algorithms (https://esbl.nhlbi.nih.gov/AbDesigner/), peptide solubility (http://pepcalc.com/), and the differential homology between SARS-CoV-2 and SARS-CoV-1, or other coronaviruses (https://blast.ncbi.nlm.nih.gov/Blast.cgi). These peptide sequences were then checked for possible negative internal amino acid interactions prior to final selection.

### Polyclonal and monoclonal antibody generation

Six-week-old female Balb/c mice (Charles River) were used for immunizations and housed at the Godley Snell Animal Facility, a Clemson University IACUC approved facility under an IACUC-approved protocol (AUP2020-0069). Mice were given a sub-cutaneous (s.c.) injection with 100 μg of immunogen in 0.9% saline containing TiterMax Gold® adjuvant on Day 1. Fourteen days later mice received a second s.c. immunization using Freund’s incomplete adjuvant. Subsequent boosters at 21-day intervals were given in saline via s.c. immunizations, and the final booster was given intraperitoneally. Five days after the last booster immunization, mice were sacrificed using slow lethal CO_2_ hypoxia, bled by cardiac puncture at the time of pneumothorax detachment to collect polyclonal anti-sera, and their spleens removed using aseptic methods. Procedures for fusion of splenocytes with Sp02–14 myelomas (ATCC, Manassas VA USA), and for screening and cloning of the resulting hybridomas have been described elsewhere [18-20]. Hybridomas were typically grown in Dulbecco’s Modified Eagle Medium (Cellgro) supplemented with 10% fetal bovine serum (FBS), 20 mM HEPES, 10 mM L-glutamine, 100 µg/mL penicillin, 100 µg/mL streptomycin, 110 µg/mL sodium pyruvate, 1% non-essential amino acids (100× stock), 4.5 g/L glucose, 10 µg/mL gentamycin and 5 µg/mL nystatin. Blood samples were allowed to clot, then centrifuged to collect overlying serum to serve as anti-sera in subsequent assays.

Primary hybridoma supernatants were then screened by ELISA against unconjugated peptide (2 μg/well) or rRBD (2 μg/well). Briefly, immunogens were diluted to 20 μg/ml in 0.01 M phosphate buffered saline, pH 7.2 (PBS) and 100 ul added to wells of a 96-well medisorp ELISA plate (Thermo Fisher Scientific #467320) and incubated overnight at 4° C. Plates contents were then discarded, and plates washed 5X with PBS containing 0.01% Tween-20 (PBST). Plate wells were then blocked overnight with 150 μl of PBS containing 10% horse serum (blocking buffer). Plate contents were then discarded, and plates received 100 μl of hybridoma supernatant, followed by incubation at room temperature for 2 hours. Plates were washed 5X with PBST, then received 100 μl goat anti-mouse IgG-AP (Thermo Fisher 1:2000 in PBS) and incubated for 2 hours at room temperature. Finally, the plates were washed 5X with PBST, then received 100 μl of *p*-nitrophenyl phosphate substrate in alkaline phosphatase buffer (100 mM NaCl, 50 mM MgCl_2_, 100 mM Tris-Cl, pH 9.5) (AP). After 30 minutes, the plates received 100 μl of stopping buffer (2 M NaOH) and were read at 405 nm and the optical density recorded.

Cells from wells with a minimal signal of three-fold optical density readings above the lowest reading wells were then cloned by limiting dilution for further testing. Hybridomas were grown to confluence and the supernatants collected by centrifugation, then treated with 0.05% NaN_3_ and stored at 4°C. Initial isotyping of the antibodies in hybridoma supernatants was carried out using Pierce Rapid Antibody Isotyping Kits for mouse (Thermo Fisher). Antibodies were subsequently used as confluent supernatants for most techniques, or further purified over protein A/G columns as needed. For obtaining purified mAbs, the hybridomas were grown in complete media as described above but containing ultralow bovine IgG serum (Fisher Scientific) to avoid purifying bovine IgG over mouse IgG. Preliminary studies demonstrated that only three hybridomas secreted antibodies specific to the immunizing peptides or rRBD. Of these, one secreted an antibody (mAb CU-P1-1) that recognizes pep-1 by ELISA, one that secreted an antibody (mAb CU-P2-20) against pep-2, and one (mAb CU28-24) that recognizes the rRBD. Immunoglobulin (Ig) genes of the three hybridomas were then sequenced (Absolute Antibody/Kerafast, Boston MA USA) using Next Generation Sequencing (NGS) to verify the presence of a single Ig and of the appropriate isotype. The unique complimentary determining region (CDR) sequences of heavy and light chains have been determined, and these Ig sequences will allow for the expression of recombinant mAbs.

### Screening polyclonal anti-sera and mAbs for reactivity against rRBD

Stored rRBD was diluted in PBS to 100 ug/ml, mixed with 6X reducing sample buffer (SB), and boiled for 7 minutes. Protein molecular weight markers (FroggoBio, Buffalo NY, USA) and 2 ug of rRBD were subjected to SDS-PAGE using 4-20% gels, then transferred to PVDF membranes. Membranes were then blocked overnight with blocking buffer, washed 3X with PBST, and then probed with the three polyclonal anti-sera samples diluted 1:500 in PBS, or the three mAbs as confluent supernatants diluted 1:3 in PBS (nominal concentration of 3 μg/ml), followed by washing 3X with PBST. The blots were then probed with goat-anti-mouse IgG-AP (Themofisher, 1:2000 in PBS) for 2 hours at room temperature, washed 3X with PBST, then developed using the substrate BCIP/NBT (Fisher Scientific) to detect the protein of interest as a dark blue band. The development process was stopped by extensive washing with tap water, the blots dried at room temperature, then imaged using a ChemiDoc™ imaging system (Biorad).

### Screening mAbs for specificity

To determine the specificity of mAbs, ELISA plates were again coated with unconjugated peptides or rRBD. Using the above-described ELISA approaches, each polyclonal anti-sera sample diluted 1:250 in PBS and mAb as confluent supernatants diluted 1:3 in PBS were used as primary antibody against peptide 1, peptide 2, or rRBD. An irrelevant mAb was used as an assay control [21]. After development, the optical density of each well was recorded at 405 nm to compare the reactivity of each mAb with each antigen.

### Immunoprecipitation of rRBD with CU28-24

From the immunoblots and ELISA data it was determined that mAb CU28-24 is specific for the native rRBD protein but does not recognize the protein under the denaturing conditions of SDS-PAGE and immunoblotting. To further validate that mAb CU-28-24 is specific for rRBD, but not recognizing either of the epitopes recognized by mAb CU-P1-1 or CU-P2-20, purified mAb CU-28-24 was applied to a protein A/G column by running the sample over the column 3X. The column was washed extensively with PBS, followed by the addition of 100 ug rRBD in 3 ml PBS which was reapplied to the column 3X. Next the column was washed 4X with 4 ml PBS, and the wash steps collected. After the final wash, proteins associated with the column were eluted with 0.8 ml aliquots of 0.05 M glycine, pH 2.5 and collected in tubes containing 0.2 ml of carbonate buffer, pH 9. These samples and wash samples were then mixed with SB and boiled, then subjected to SDS-PAGE and immunoblotting. One lane of the gel contained 2 ug of rRBD as a positive control. The blots were blocked overnight with PBS containing 10% horse serum, washed 3X, then probed with FITC conjugated [22] purified mAb CU-P2-20 diluted in PBS to 5 μg/ml. The blot was washed 3X, dried, and imaged for fluorescent banding using a ChemiDoc™ imaging system (Biorad).

### SARS-CoV-2 surrogate virus neutralization assays using antisera

A commercially available kit to quantify the ability of the three pAbs to neutralize the binding of RBD to ACE-2 was obtained from GenScript, Piscataway NJ (kit #L00847). The methods followed those provided by the supplier. The wells were precoated by the manufacturer with rACE-2 protein. In brief, pAb samples were diluted 1:10, 1:50, 1:100, and 1:200 in assay dilution buffer in the provided assay ELISA plate. Diluted samples and HRP-labeled RBD were mixed, incubated as directed, and then applied to the ACE-2-coated plate in duplicates. After 30 minutes incubation, the plates were washed extensively with provided wash buffer, and HRP substrate was added. After 30 minutes the reaction was stopped with provided stop buffer and the optical density at 450 nm was recorded and the data was processed as directed by the manufacturer and expressed as percent signal inhibition. Positive and negative control anti-sera solution were provided by the manufacturer.

### Plaque reduction neutralization test (PRNT) assays using mAbs

Virus neutralization assays were carried out at Georgia State University under the supervision and generosity of Dr. Mukesh Kumar using a certified Biosafety Level 3 laboratory as described previously [23]. The details of PRNT follow standard protocols [24]. The three mouse mAbs were diluted to 50 μg/ml in PBS to begin the test. Vero E6 cells were used as the host target model and live SARS-CoV-2 strain USA-WA1/2020 virus was used for infection. The antibodies were diluted serially from 1:4 to 1:4096 and PRNT was conducted using Wuhan (B.1) strain of SARS-CoV-2. The highest dilution of serum resulting in 50% reduction in the number of plaques compared to the growth of the virus control was determined.

### Mouse infection and tissue harvesting

Brain and lung tissues from infected and control mice were a generous gift from Dr. Mukesh Kumar, Georgia State University. Hemizygous K18-hACE2 mice were purchased from the Jackson Laboratory (Bar Harbor, ME). All the animal experiments were conducted in a certified Animal Biosafety Level 3 (ABSL-3) laboratory at Georgia State University (GSU). The protocol was approved by the GSU Institutional Animal Care and Use committee (Protocol number A20044). Six-week-old hemizygous K18-hACE2 mice were infected with 10^5^ plaque-forming units (PFU) of SARS-CoV-2 strain USA-WA1/2020 under ABSL-3 containment by intranasal inoculation. Animals in the control group received equivalent amounts of sterile PBS via the same route. Roughly equal numbers of male and female mice were used. Animals were weighed and their appetite, activity, breathing and neurological signs assessed twice daily. On day 5 post infection, the animals were anesthetized and perfused with 10 mL PBS followed by 10 mL 4% paraformaldehyde (PFA). The tissues were collected and stored in PFA.

### Tissue Immunohistochemistry

Lungs and brains were grossed, recorded, and then were placed in 10% buffered formalin. After three days, the formalin was removed and replaced with 70% ethanol for storage. Tissues were processed and embedded in ParaPlast Extra by the Louisiana Animal Disease Diagnostic Laboratory (LADDL) at Louisiana State University School of Veterinary Medicine. Newly embedded tissues were then sectioned and affixed to charged slides for H&E staining and immunohistochemistry. For antigen retrieval, tissue slices were microwave-heated in either pH 6 citrate buffer or pH 9 Tris-EDTA. Tissues were then encircled with a Liquid Blocker Super mini pen to isolate tissue slices, thus dividing each slide into two sections to allow one section to act as a reagent control by withholding primary antibody treatment. Additional screening assays included isotype controls. All sections were subsequently blocked with 5% horse serum (VectaStain®) blocking buffer for 17 minutes. Next, the antibody, as hybridoma supernatant diluted 1:3 in PBS with 1.5% horse serum, was added dropwise to the slides and incubated overnight at 4 °C. After incubation, the slides were washed prior to addition of the secondary antibody (goat anti-mouse IgG-Alexafluor488, 5 ug/ml, Invitrogen) and incubated for 2 hours at room temperature. Slides were washed 5X in PBST, stained with DAPI for 5 min, washed again. Finally, slides were mounted, cover-slipped and imaged.

### Screening polyclonal anti-sera and mAbs for reactivity against Omicron BA.2 and BA.4.5 variant rRBDs

Recombinant Omicron RBD proteins from variants BA.2 and BA.4.5 were obtained from Fisher Scientific. To determine the specificity of mAbs, ELISA plates were coated with rRBD from the two variants as well as from the original Wuhan strain. Using the above-described ELISA approaches, each mAb as confluent supernatants diluted 1:3 in PBS were used as primary antibody. After development, the optical density of each well was recorded at 405 nm to compare reactivity of each mAb with each antigen.

## RESULTS

### Immunizations and resulting anti-sera

In this study, mice were immunized with two different synthetic peptides mapping to the RBD region of the SARS-CoV-2 spike protein. Mice were also immunized with commercially available rRBD expressed in HEK cells. The product is approximately 37 kDa (Figure 1). This product, like most recombinant proteins, tends to degrade slightly upon thawing and refreezing.

**Figure 1.**
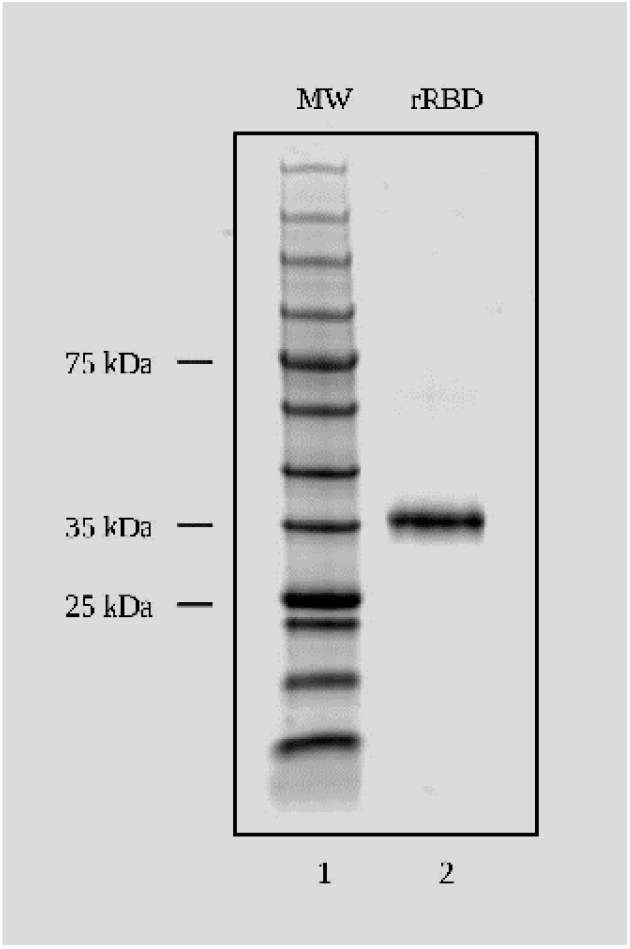
SDS-PAGE and Coomassie blue staining of SARS-CoV-2 (Wuhan) recombinant RBD used for immunizations and antibody screening. SDS-Page and Coomassie blue staining of rRBD protein. Lane 1: 12 ul protein molecular weight marker. Lane 2: 2 μl of 37 kDa rRBD protein.

Serum from mice immunized with peptides and whole rRBD was used as a source of anti-sera to determine if mice responded immunologically to the immunogen associated with RBD. Anti-sera from all three immunogens recognized rRBD by Western blots (Figure 2). In each case, the partially degraded rRBD product was also recognized by the respective anti-sera. Anti-sera were also used to determine the degree of specificity against the respective peptide and rRBD by ELISA. Anti-sera from mice immunized with peptide 1 (P1) recognized P1, but not P2, and recognized whole rRBD (Figure 3). Likewise, anti-sera from mice immunized with peptide 2 (P2) recognized P2, but not P1, and recognized whole rRBD. Of note, neither anti-sera from P1- or P2-immunized mice were as reactive against rRBD as anti-sera from rRBD-immunized mice.

**Figure 2.**
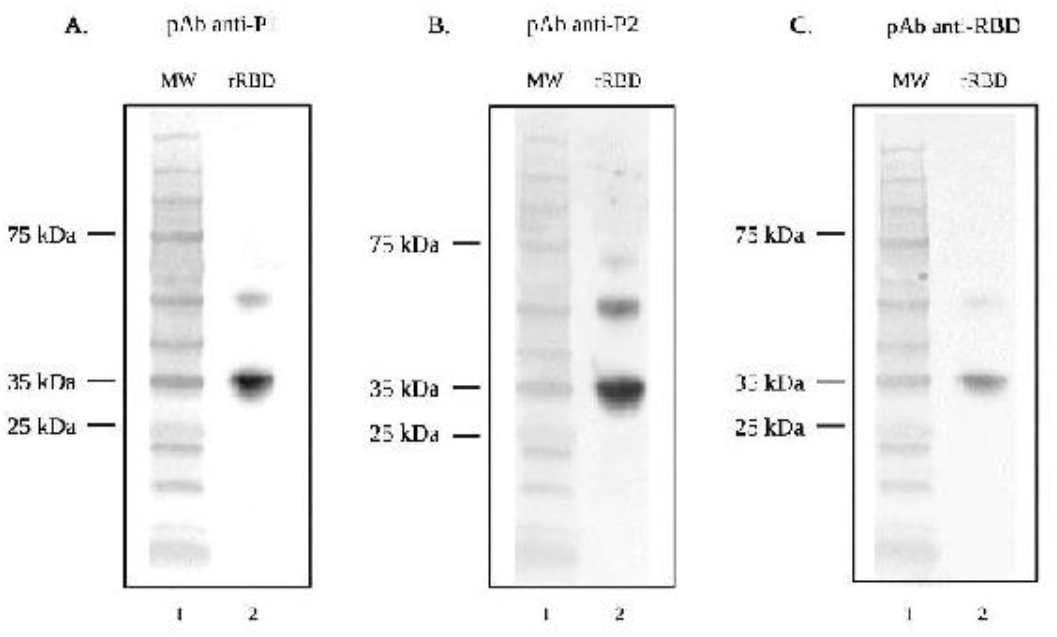
SDS-Page and Western blot analysis demonstrating recognition of SARS-CoV-2 (Wuhan) rRBD by anti-sera. Lane one for each Western blot contains the protein molecular weight marker. Lane two of each Western blot contains rRBD (37 kDa) probed with (A) pAb anti-P1, (B) pAb anti-P2, and (C) pAb anti-RBD followed by goat anti-mouse IgG-AP secondary antibody.

**Figure 3.**
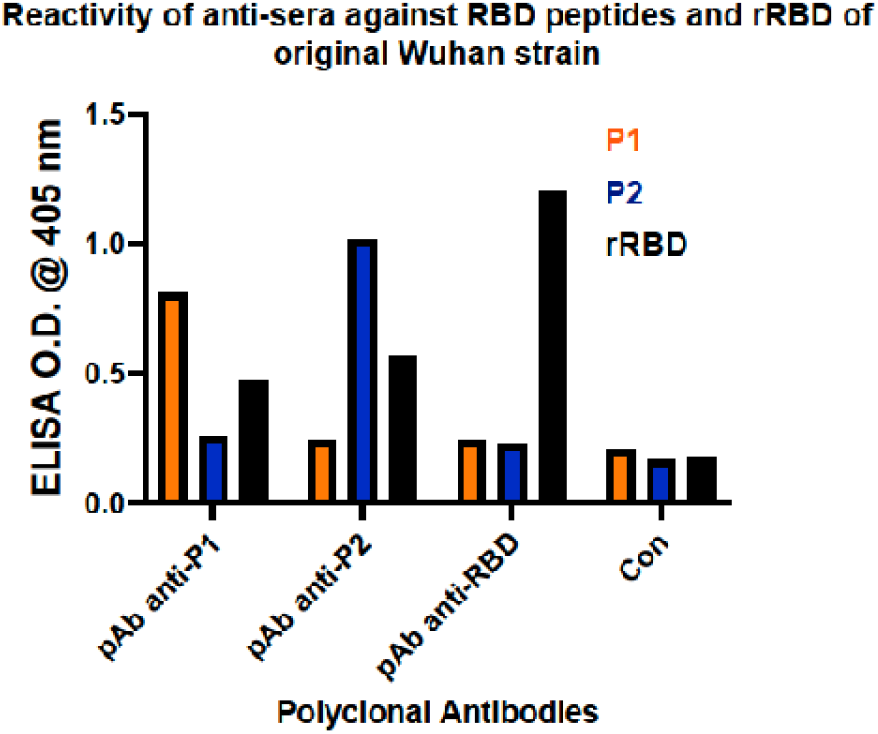
Enzyme-linked immunosorbent assay (ELISA) for reactivity of anti-sera against RBD peptides and rRBD of SARS-CoV-2 (Wuhan) Reactivity of anti-sera from mice immunized against P1, P2, and rRBD against whole rRBD and respective peptides.

### Surrogate viral neutralization assay with anti-sera

The three anti-sera samples from immunizations were used to determine if they contained antibodies that could neutralize the ability of RBD binding to ACE-2 using a commercially available surrogate viral neutralization assay kit. In this assay, HRP-labeled RBD is incubated with dilutions of anti-sera, and this mixture is added to plates coated with ACE-2. Data is recorded as percent signal inhibition. Only anti-sera from rRBD mice inhibited the binding of RBD to ACE-2, and this ability was at the maximal level detected by the kit (Figure 4).

**Figure 4.**
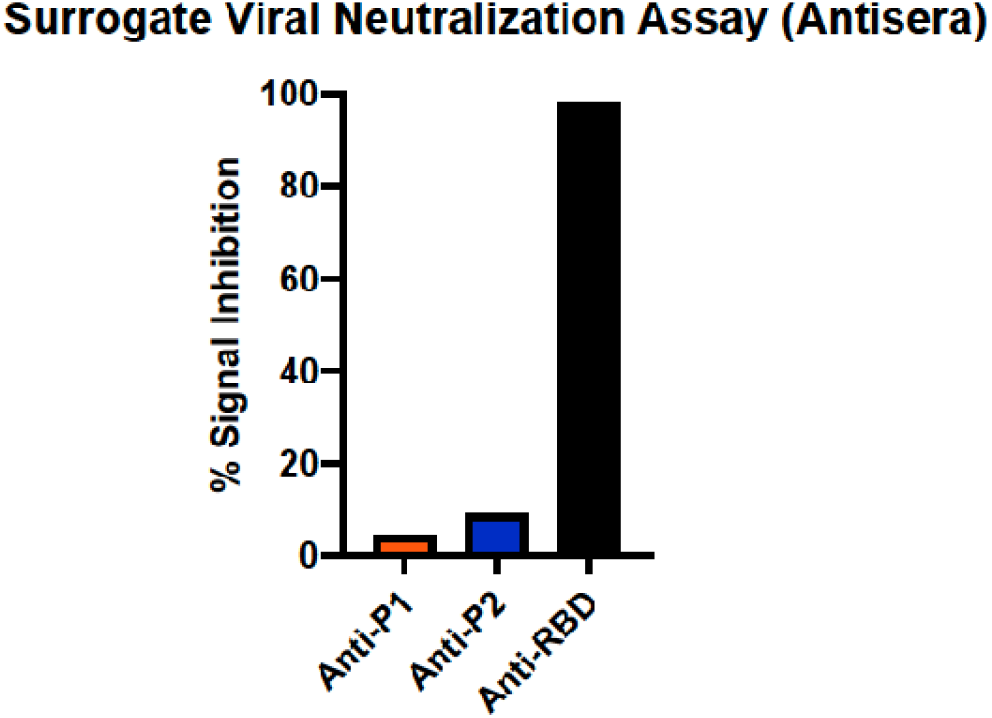
Surrogate Viral Neutralization Test (sVNT) demonstrating neutralization of RBD-ACE2 binding by anti-sera. Neutralization activity of anti-sera represented by percent signal inhibition of HRP-labeled RBD binding to ACE-2 by anti-sera from mice immunized against peptides P1, P2, and whole rRBD.

### Characterization of monoclonal antibodies

From immunizations and steps to develop cloned hybridomas secreting mAbs against P1, P2, and rRBD, three mAbs were cloned, isotyped, and named CU-P1-1 (IgG_1_ κ) against P1, CU-P2-20 (IgG_1_ κ) against P2, and CU-28-24 (IgG_2b_ κ) against rRBD. Using an ELISA approach to determine specificity, CU-P1-1 recognizes P1, but not P2, and slightly reacts with rRBD (Figure 5). CU-P2-20 recognizes the P2 peptide, but not P1, and is highly reactive against rRBD. CU-28-24 does not recognize either P1 or P2 but does react highly with rRBD. These hybridomas were then submitted to Kerafast, Inc. for sequencing to verify only one immunoglobulin gene (H + L) is expressed and the correct isotype. Sequencing also provided the sequences for each of the six CDR regions responsible for epitope binding and are protected by non-disclosure agreements with Clemson University Research Foundation (CURF).

**Figure 5.**
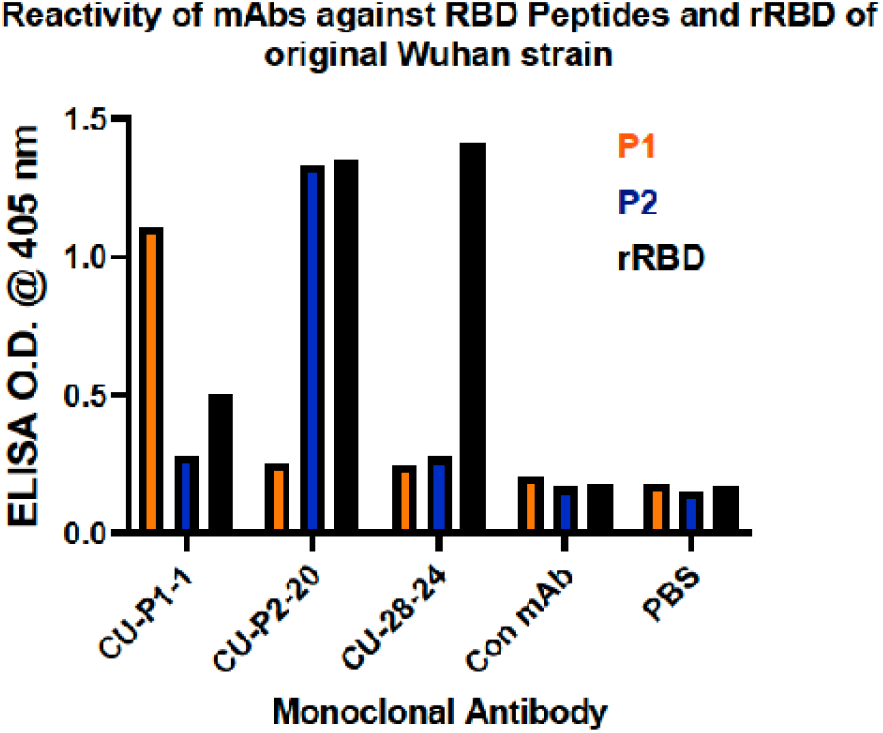
Enzyme-linked immunosorbent assay (ELISA) for reactivity of mAbs against RBD peptides and rRBD of SARS-CoV-2 (Wuhan). Reactivity of mAbs CU-P1-1, CU-P2-20, and CU-28-24 against P1, P2, and whole rRBD of original Wuhan strain of SARS-CoV-2

Each of the three mAbs were then tested by immunoblotting to verify the recognition of rRBD at the expected molecular weight. Monoclonal antibodies CU-P1-1 and CU-P2-20 recognize rRBD and recognize slight degradation products of the protein (Figure 6). However, mAb CU-28-24 does not recognize rRBD by immunoblotting, which is likely due to epitope destruction under the denaturing conditions of SDS-PAGE.

**Figure 6.**
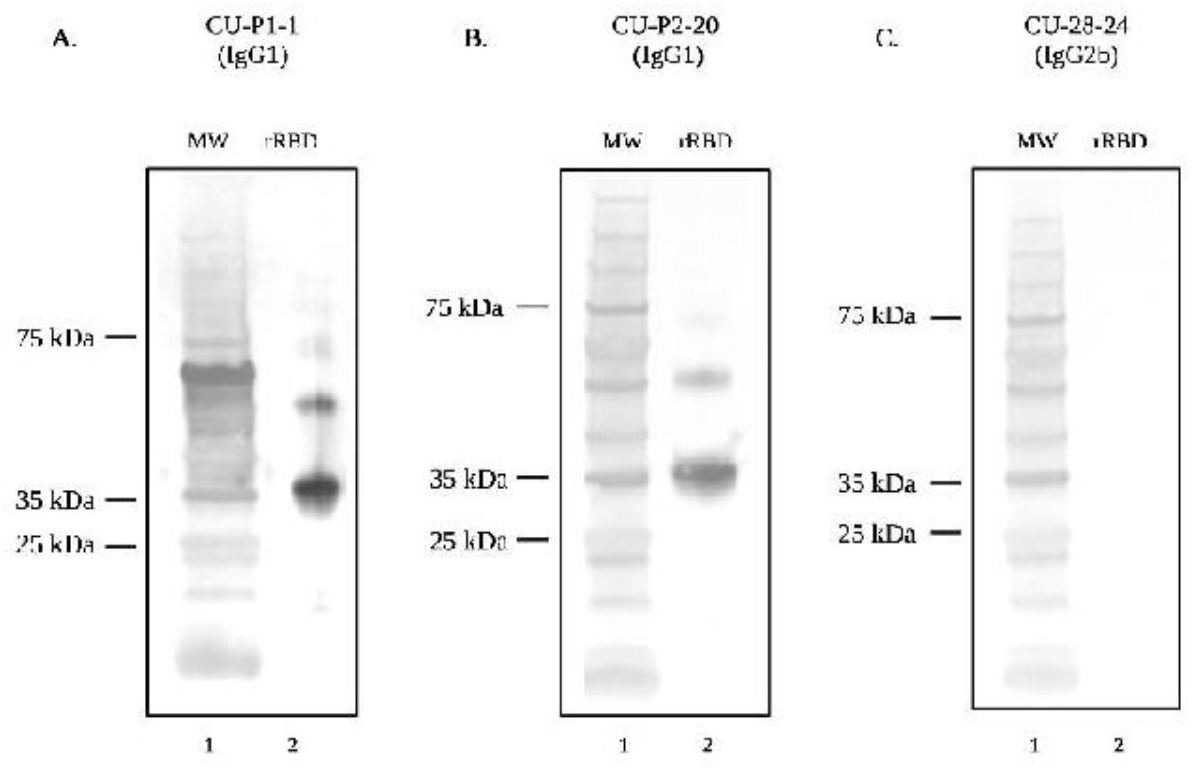
SDS-PAGE and Western blot analysis demonstrating recognition of rRBD of SARS-CoV-2 (Wuhan) by mAbs. Lane one for each Western blot contains the protein molecular weight marker. Lane two of each Western blot contains rRBD (Wuhan) probed with (A) mAb CU-P1-1, (B) mAb CU-P2-20, and (C) mAb CU-28-24 followed by goat anti-mouse IgG-AP secondary antibody.

### Immunoprecipitation of rRBD with CU-28-24 and detection with CU-P2-20

Efforts to immunoprecipitate rRBD with Protein-A/G bound CU-28-24 were successful in that after multiple washings of the bound column, rRBD was eluted from the column (Figure 7). FITC labeled CU-28-24 was then used to detect both eluted rRBD and pure rRBD by immunoblotting.

**Figure 7.**
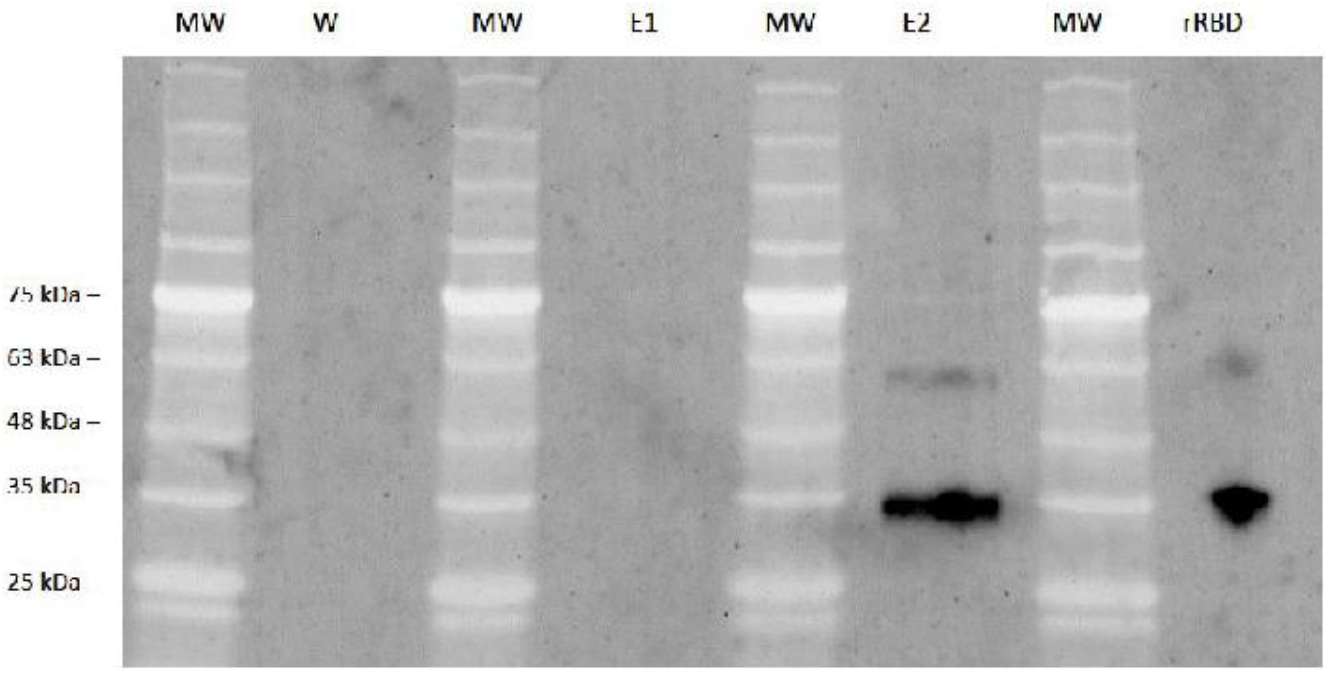
Immunoprecipitation of rRBD with mAb CU-28-24 and subsequent immunoblotting with mAb CU-P2-20. Purified mAb CU-28-24 was captured with protein A/G column, washed extensively, then rRBD was applied to the column to bind rRBD with mAb CU-28-24. After additional wash steps, the contents of the column were eluted in 1 ml fractions. The last wash step and first two elutions were subjected to SDS-PAGE and immunoblotting with FITC-labeled mAb CU-P2-20 and imaged. MW: molecular weight markers. rRBD: commercially sources rRBD as an internal control. Both precipitated and control rRBD were labeled.

### Viral neutralization test with monoclonal antibodies

The plaque reduction neutralization test is the gold standard for determining the neutralization titer of antibodies against live virus. Under the conditions carried out in the BSL-3 facility at Georgia State University, CU-P1-1 had no ability to block the virus from entering host Vero E6 cells, and CU-P2-20 neutralized 50% of the virus with only a small titer of 4 (Figure 8). However, CU-28-24 had a PRNT_50_ titer of 256.

**Figure 8.**
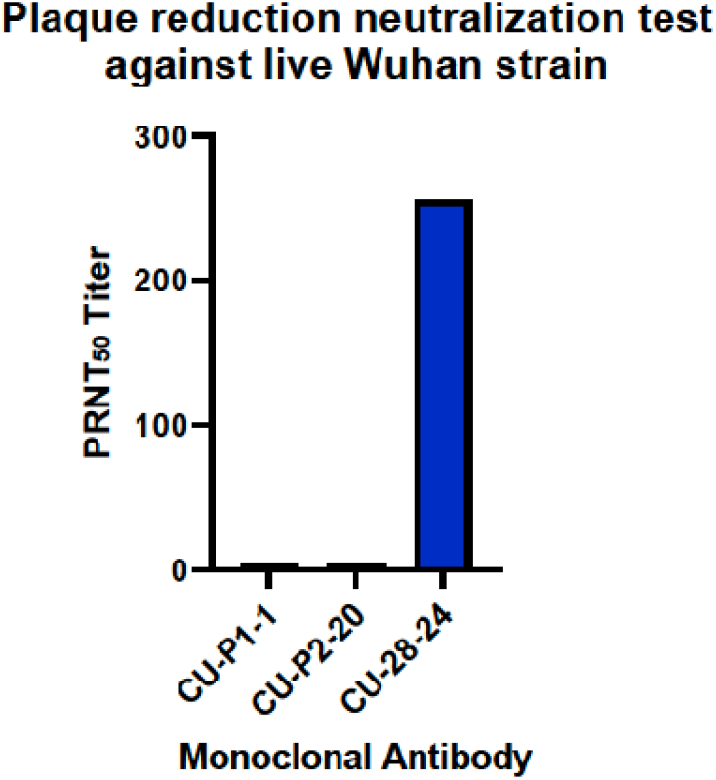
Plaque Reduction Neutralization Test of mAbs against live SARS-CoV-2 (Wuhan). Plaque Reduction Neutralization Test_50_ titer of mAbs CU-P1-1, CU-P2-20, and CU-28-24 against live SARS-CoV-2 virus (Wuhan).

### Immunohistochemistry using CU-P1-1, CU-P2-20, and CU-28-24

Mouse tissues infected with SARS-CoV-2 (Wuhan) show intensive staining with mAbs CU-P2-20 and CU-28-24 (Figure 9). Alternatively, CU-P1-1 was only marginal in recognizing the virus in lung and brains, despite efforts to optimize conditions (images not shown). It was noted throughout optimization steps that antigen retrieval required a buffer of pH 9 for mAb CU-P2-20 and pH 6 for mAb CU-28-24.

**Figure 9.**
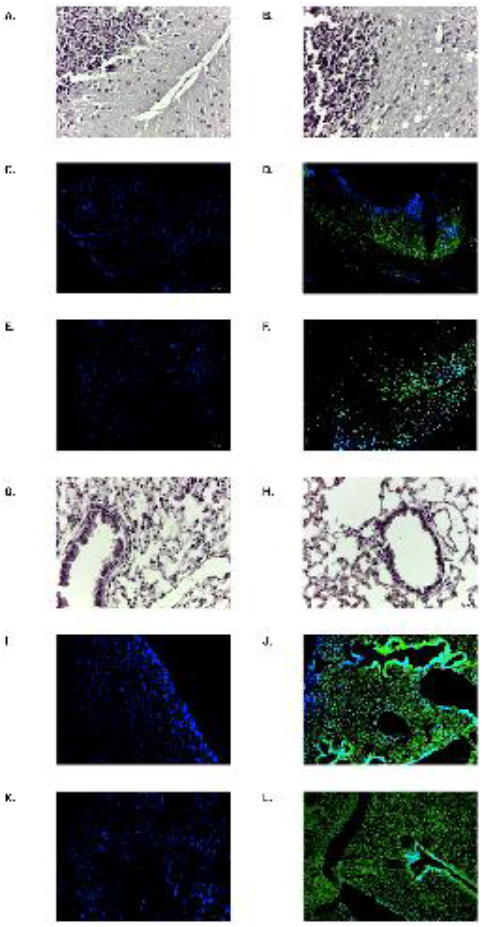
H&E staining and immunohistochemistry results of SAR-CoV-2 infected tissues with mAb CU-P2-20 and mAb CU-28-24. Tissues from either control or infected brains or lungs from SARS-CoV-2 (Wuhan strain) were H&E stained or were probed with mAbs, washed, then probed with FITC-labeled goat anti-mouse IgG and DAPI, washed and cover-slipped prior to imaging. (A) H&E staining of non-infected Brain (B) H&E staining of infected brain. (C) Non-infected brain: mab CU-28-24. (D) Infected brain: mAb CU-28-24. (E) Non-Infected Brain: mAb CU-P2-20. (F) Infected Brain: mAb P2-20. (G) H&E staining of non-infected lung. (H) H&E staining of infected lung. (I) Non-Infected Lung: mAb CU-28-24. (J) Infected lung: mAb CU-28-24. (K) Non-Infected Lung: mAb CU-P2-20. (L) Infected Lung: mAb CU-P2-20

### Reactivity of mAbs against RBD of recent Omicron strains BA.2 and BA.4.5

Recombinant RBD derived from Omicron strains BA.2 and BA.4.5. were used to coat ELISA plates and then probed with the three mAbs to compare reactivity with rRBD from the original Wuhan strain. CU-P1-1 and CU-P2-20 demonstrated relatively low reactivity against the two strains of Omicron tested in this study (Figure 10). As expected, each of these two were reactive against the rRBD from the Wuhan strain. CU-28-24, however, strongly recognized BA.2 and BA.4.5. rRBDs, as well as rRBD derived from the Wuhan strain.

**Figure 10.**
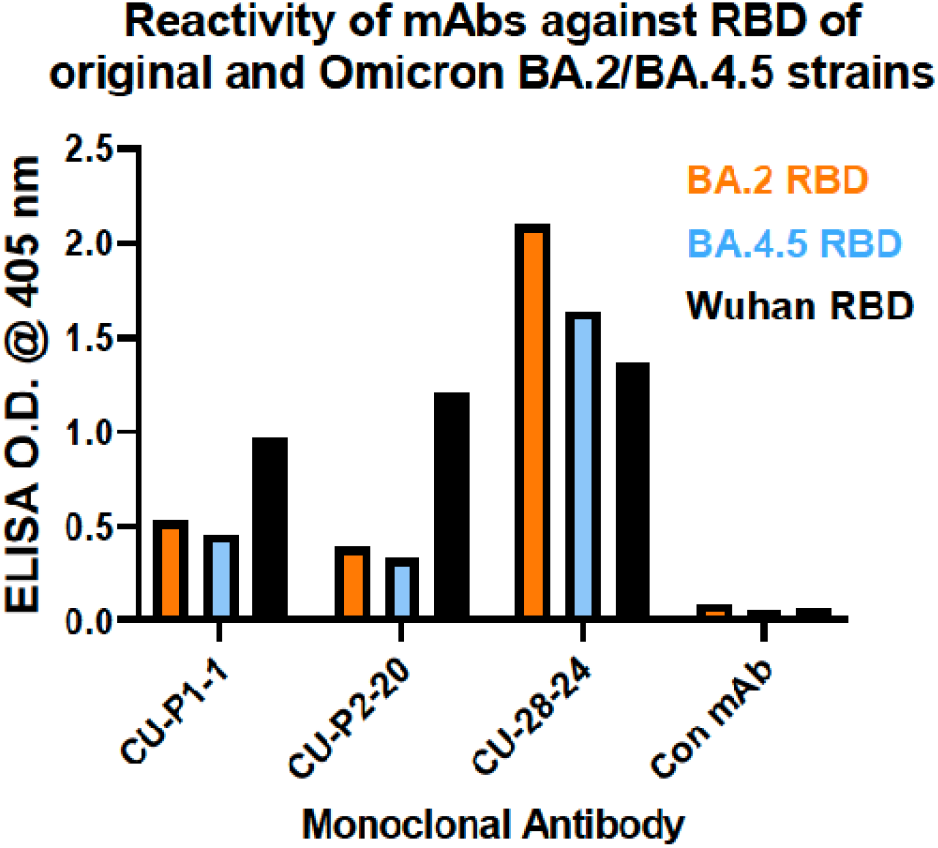
Enzyme-linked immunosorbent assay (ELISA) of reactivity of mAbs against rRBDs derived from SARS-CoV-2 (Wuhan) and Omicron variants BA.2 and BA.4.5. Reactivity of mAbs CU-P1-1, CU-P2-20, and CU-28-24 against RBD of original Wuhan strain and Omicron BA.2 and BA.4.5 variant strain RBDs.

## DISCUSSION

In this study, a panel of three different monoclonal antibodies against non-overlapping epitopes within the RBD of the original Wuhan SARS-CoV-2 virus were successfully generated and characterized. Two of the three epitopes recognized by the panel are defined by the specific synthetic peptide (peptide 1 vs peptide 2) chosen for immunization, while the third antibody (CU-28-24) recognizes a yet to be determined epitope outside of the region of the two peptide sequences. The most common means to determine the epitope of mAb CU-28-24 would be peptide walking, in which overlapping synthetic peptides are generated to then screen the antibody by ELISA. Likewise, the peptide recognized by ELISA during the “walking” should also block the binding of CU-28-24 to whole rRBD in both ELISA and immunoblotting techniques. Based on the quality of mAb CU-28-24 as described herein, future studies should be carried out to determine the specific epitope.

One of several observations from this study is that mAb CU-P2-20 reacts with P2 and rRBD equally well, but mAb CU-P1-1 against P1 does not bind well to rRBD in ELISAs. This observation may indicate that the region or epitope associated with P1 is not as immunogenic as predicted or is simply not available to the antibody in native full rRBD. Though Hopps & Woods hydrophilicity plotting and NIH Abdesign programs predict P1 to be highly immunogenic, there may have been negative interactions between amino acids within the peptide that prevented the epitope to be fully expressed. For example, the original peptide designed by NIH Abdesign was AWNSNNLDSKVGGNYNYLYR, but this peptide is completely insoluble in water or PBS for immunizations. To make this more soluble and suitable for immunization, the peptide was shortened to NSNNLDSKVGGNYNY prior to cysteine addition on the N-terminus for KLH conjugation. However, this sequence leads with asparagine (N) at the N-terminus, there are multiple N within the sequence, and two adjacent internal glycines (G), leading to the likelihood of hampered confirmational structure compared to the native rRBD [25]. Of note, mAb CU-28-24 did not recognize rRBD by SDS-PAGE/immunoblotting under reducing conditions, suggesting that the epitope within RBD may have been more readily revealed under these conditions. On the other hand, P2 (<u>QTGKIADYNYKLPDDFTG)</u> which was conjugated with KLH at the C-terminus using an additional cysteine and thus did not exhibit such predictable issues, was water soluble. Monoclonal antibody CU-P2-20, like mAb CU-P1-1, readily recognizes RBD in SDS-PAGE/immunoblotting and together with characteristics in ELISA mAb CU-P2-20 should have applications in a variety of endpoints. Of note, based on the amino acid sequence of the RBDs of SARS-CoV and SARS-CoV-2, mAb CU-P1-1 may distinguish between the two and mAb CU-P2-20 should have activity against both.

The observation that mAb CU-28-24 recognizes RBD by ELISA, but not by SDS-PAGE/immunoblotting indicates that its specific epitope is destroyed during the denaturing conditions. However, this does not hinder the ability of the antibody to immunoprecipitate rRBD for recognition by CU-P2-2-, nor does it hinder the ability of the antibody to neutralize the live virus in PRNT assays. Therefore, this antibody should be of high value in future studies with other variants of concern. To that end, mAb CU-28-24 readily recognizes rRBDs from the Omicron BA.2 variant and the BA.4/BA.5 variant, and the degree of response in ELISA was comparable to activity against rRBD from the original Wuhan strain. Future studies with CU-28-24 should be conducted to determine if current variants of concern and those appearing in the future are neutralized by this antibody. In this regard, mAb CU-P1-1 and mAb CU-P2-20 have no value in terms of viral neutralization.

As previously addressed, anti-viral antibodies may have several functions outside of the ability to neutralize a virus of concern. One endpoint of high value with mAbs is in immunohistochemistry for detecting the virus in specific tissues. Using IHC and immunofluorescent labeled secondary antibody, it seems that CU-P1-1 does not work well for IHC. However, mAbs CU-P2-20 and CU-28-24 show viral infection in infected lungs and brains of mice exposed to the virus, and this activity seems to be concentrated in focal regions, consistent with observations made by others [26-28]. While some studies use antibodies generated against nucleocapsid proteins [29,30], many others use antibodies against the RBD [31]. Of concern would be the high rate of mutation within the RBD of variants of concern post emergence of the Wuhan strain, making some original RBD-based antibodies ineffective. However, based on the ELISA results against Omicron BA.2 and BA.4.4, mAb CU-28-24 should be a good reagent going forward.

The original hypothesis was that defined synthetic peptides derived from the RBD region of the SARS-CoV-2 virus would be a viable approach to developing high quality antibodies with a broad range of applications. Considering the importance of neutralizing antibodies for possible therapeutic endpoints, this seems not to be the case, and perhaps larger proteins for immunization would be best, as demonstrated by mAb CU-28-24. However, a panel of only three mAbs from this study is not enough to make broad statements about the utility of peptide-based immunogens compared to larger, more native proteins from which the peptide sequences were derived. Regardless, this study provides at least two very good mAbs for continued research related to COVID-19 and the emerging variants of concern. Considering the high degree of cross-reactivity of mAb CU-28-24 with Omicron variants BA.2 and BA.4.5, it may be useful in clinical settings, especially if the antibody were to be humanized. In this case, the mouse nucleic acids corresponding to the six CDR regions would be grafted into a human IgG immunoglobulin molecule and expressed as a recombinant protein in CHO cells.

## CONCLUSION

Despite the successes with the development of vaccines, anti-viral drugs, and monoclonal antibodies for the treatment of SARS-CoV-2 and its many evolved variants of concern, there is still a need for high-quality monoclonal antibodies. This is especially true for variants of concern, such as the current Omicron variants, that are resistant to earlier monoclonals targeted against the original Wuhan strain. Going forward, mAb CU-P2-20 and mAb CU-28-24, may have both research and clinical applications. These hybridomas have been sequenced, thereby allowing scientists to express these valuable mAbs as recombinant proteins. This will supplant the need for careful and laborious hybridoma maintenance going forward. The synthetic peptides and hybridomas/mAbs are under the intellectual property management of the Clemson University Research Foundation, and the three CDRs have been submitted as an invention disclosure for further patenting and commercialization.

## Acknowledgements

A sincere appreciation is extended to doctoral student Alyssa Whisel for help in maintaining hybridomas at Clemson University (CU). This work was funded by a Clemson University Research Foundation (CURF) technology maturation fund (Tech ID: 2020-064) and by the Georgia State University (GSU) Institutional funds. We thank members of the GSU High Containment Core and the Department for Animal Research for assistance with the experiments. We also thank Absolute Antibody/Kerafast (Boston MA USA) for providing immunoglobulin gene sequences.

